# Discovery of a first-in-class small molecule ligand for WDR91 using DNA-encoded chemical library selection followed by machine learning

**DOI:** 10.1101/2023.08.21.552681

**Authors:** Shabbir Ahmad, Jin Xu, Jianwen A Feng, Ashley Hutchinson, Hong Zeng, Pegah Ghiabi, Aiping Dong, Paolo A Centrella, Matthew A Clark, Marie-Aude Guié, John P Guilinger, Anthony D Keefe, Ying Zhang, Thomas Cerruti, John W. Cuozzo, Moritz von Rechenberg, Albina Bolotokova, Yanjun Li, Peter Loppnau, Alma Seitova, Yen-Yen Li, Vijayaratnam Santhakumar, Peter J. Brown, Suzanne Ackloo, Levon Halabelian

## Abstract

WD40 repeat-containing protein 91 regulates endosomal phosphatidylinositol 3-phosphate levels at the critical stage of endosome maturation and plays vital roles in endosome fusion, recycling, and transport by mediating protein-protein interactions. Due to its various roles in endocytic pathways, WDR91 has recently been identified as a potential host factor responsible for viral infection. We employed DNA-Encoded Chemical Library (DEL) selection against the WDR domain of WDR91, followed by machine learning to generate a model that was then used to predict ligands from the synthetically accessible Enamine REAL database. Screening of predicted compounds enabled us to identify the hit compound **1**, which binds selectively to WDR91 with a K_D_ of 6 ± 2 μM by surface plasmon resonance. The co-crystal structure confirmed the binding of **1** to the WDR91 side pocket, in proximity to cysteine 487. Machine learning-assisted structure activity relationship-by-catalog validated the chemotype of **1** and led to the discovery of covalent analogs **18** and **19**. Intact mass LC-MS and differential scanning fluorimetry confirmed the formation of a covalent adduct, and thermal stabilization, respectively. The discovery of **1, 18, 19**, accompanying SAR, and co-crystal structures will provide valuable insights for designing more potent and selective compounds against WDR91, thus accelerating the development of novel chemical tools to evaluate the therapeutic potential of WDR91 in disease.

## Introduction

Eukaryotic cells depend on a finely controlled endo-lysosomal system to regulate the protein and lipid content of the plasma membrane, maintain intracellular quality control, and signaling ^1^. Endosomal maturation is controlled by the Rab family of small GTPases activity, phosphoinositides, and series of effector proteins^2^. Effective switching from Rab5 to Rab7 as well as controlling the levels of phosphatidylinositol 3-phosphate (PtdIns3P) in early-endosomes is crucial for early-to-late endosome maturation and recycling^3^. PtdIns3P is generated in endosomes by the activity of class III PI3K complex, which is composed of Vps34, p150, and Beclin1^4^, and downregulated via dephosphorylation by the activity of myotubularin family of 3-phosphatases^5^.

Normal endosome maturation is required for key physiological processes including neuronal development^6^ and studying changes in the endo-lysosomal system in relation to cellular physiology is a fundamental step in elucidating human pathologies ^7^. By using genome-wide CRISPR screens, Burkard et al. and others have recently identified a suite of host cell factors, including WDR91 and WDR81, that promote viral infection and replication ^8–10^. Viral entry to host cells via the endosome is well documented ^11,12^, however it is unclear whether the role of WDR91 involves trafficking and degradation of viral restriction factor tetherin ^13^ and/or differential phosphorylation^8^.

WD40 repeat-containing protein 91 (WDR91) is an 83 kDa protein composed of a coiled-coil motif (aa 183-215), a disordered region (aa 265-391) followed by a C-terminal WD40 repeat (WDR) domain (aa 392-747), which was reported to play a role in endosome maturation, retromer-mediated endosomal cargo recycling^14^, and lysosome fusion^15^. By interacting with WDR81, Beclin1^2^ and Rab7 (GTP-bound), WDR91 suppresses the class III PI3K complex activity, thus downregulating the PtdIns3P levels in endosomes ^1,2,6,13–16^.

WDR91 belongs to a large family of WDRs with a characteristic β-propeller fold (Fig 3A). WDRs are involved in key cellular processes including cell division, transcription and epigenetic regulations, DNA-damage sensing, and repair ^17,18^. Several members of the WDR protein family were shown to be druggable, including WDR5 ^19,20^, EED ^18^ and, recently, DCAF1 ^21,22^. The hit-finding strategies ranged from HTS to the more novel DNA-Encoded Chemical Library (DEL) selection followed by machine learning (ML) and use of the resultant model to predict active virtual catalog compounds ^21,23^ .

To better understand the role of WDR91 in physiology and viral infection, we aimed to develop small molecule chemical probes targeting the WDR domain of WDR91. Here, we describe the discovery of first-in-class small molecule ligands of WDR91 by using DEL selection followed by machine learning (ML) and the ordering of predicted catalog compounds ^21,23^. Primary screening and hit confirmation were performed by a suite of biophysical techniques, such as, surface plasmon resonance (SPR), differential scanning fluorimetry (DSF), mass spectrometry, and X-ray crystallography.

## Results and Discussion

### Screening and orthogonal confirmation of DEL-ML-predicted ligands

To identify the initial hits, the WDR domain of WDR91(residue range 392-747), hereafter referred to as ctdWDR91, was screened against the X-Chem DEL deck using affinity-mediated selection. This encoded compound collection comprises over 125 billion different small molecules which were synthesized using a wide range of different chemistries across multiple individual DELs using a split-and-mix method^24^. A graph convolutional neural network (GCNN) ML model, trained with affinity-mediated selection data, predicted 200 diverse putative drug-like ligands from the Enamine REAL and MCule databases. Of these, 150 compounds were successfully synthesized (**Table S1**).

Validation of the DEL-ML predicted compounds was performed by SPR. A primary screen (at 45 μM) by SPR was conducted using biotinylated ctdWDR91, and the well-studied WDR5 was used as a negative control ^25^. Eleven compounds showed specific binding to ctdWDR91 (**Fig. 1B**). The most potent compound (**1**) featured 75% binding (with respect to the maximum theoretical binding capacity) specifically to ctdWDR91 and was confirmed in a dose-response titration yielding *K*_D_ of 6 ± 2 μM (**Fig. 1C, 1D**). The affinity was also confirmed from a re-purchased solid sample.

**Figure 1:**
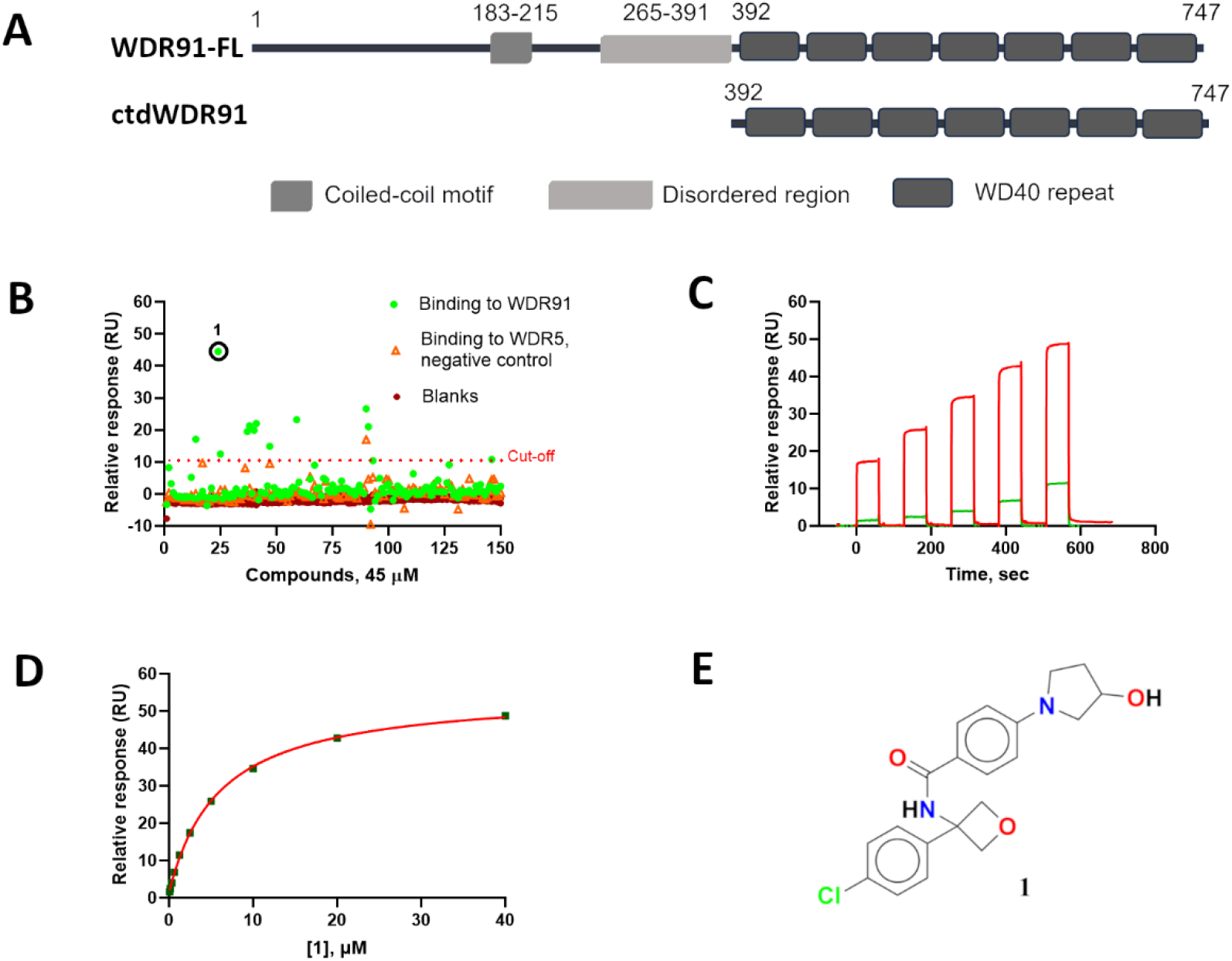
Screening and hit validation by SPR. (A) Schematic representation of full-length WDR91 (WDR91-FL) and truncated C-terminal WDR domain (ctdWDR91). (**B**) Single-dose SPR screening of predicted compounds against ctdWDR91 (residue range 392-747) identified **1** (black circle) as a hit compound. (**C**) Validation by dose-response titration of **1** against ctdWDR91 in SPR. Raw sensorgram data for concentrations ranging from 0.078 - 1.25 μM (green) and 2.5 – 40 μM (red). The 10-point titration was performed with two-fold dilution in triplicates. (**D**) Response vs concentration plot for the hit compound **1**. The *K*_D_ of 6 ± 2 μM was obtained by curve-fitting. One replicate is shown here for clarity. (**E**) Chemical structure of compound **1**.

DEL-ML-driven hit-expansion predicted 84 compounds, of which 27 exhibited binding at a single concentration (45 μM) screening by SPR. Follow-up dose-response titration of these 27 compounds confirmed binding for several with *K*_D_ values between 6-100 μM, while the other compounds either displayed low % binding (% R_max_) or exhibited poor solubility at higher compound concentrations (**Table S2**). Several analogs showed comparable binding as **1** to ctdWDR91, with *K*_D_ values ranging from 6-15 μM **(Fig. 2**). While none showed higher-affinity binding than **1**, some SAR was evident. A wide variety of substituted amides were tolerated (**Fig. 2**). Simple amides with heteroalkyl substituted phenyl group (**1, 2, 6, 11, 13, 15, 16**) as well as a wide variety of heterocycles were tolerated (**3-5, 7, 8, 12, 14**). Of the variations tested, an oxetane central ring was optimal **(1-8)**. The corresponding analogues with cyclopropyl group (**11**), hydroxyethyl group (**10, 13, 16**), methyl group (**14, 15**) and no substitutions (**9, 12**), were also tolerated, suggesting that other substitutions could further improve binding. At the oxetane ring, close analogue of the optimal p-chlorophenyl group (**1-8, 11, 13, 16**), m-chloro (**10, 14**) as well as other heterocyclic substitutions (**9, 12**) are also tolerated which indicates that other substitutions at this position could also be tolerated and could be used to design more potent ligands.

**Figure 2:**
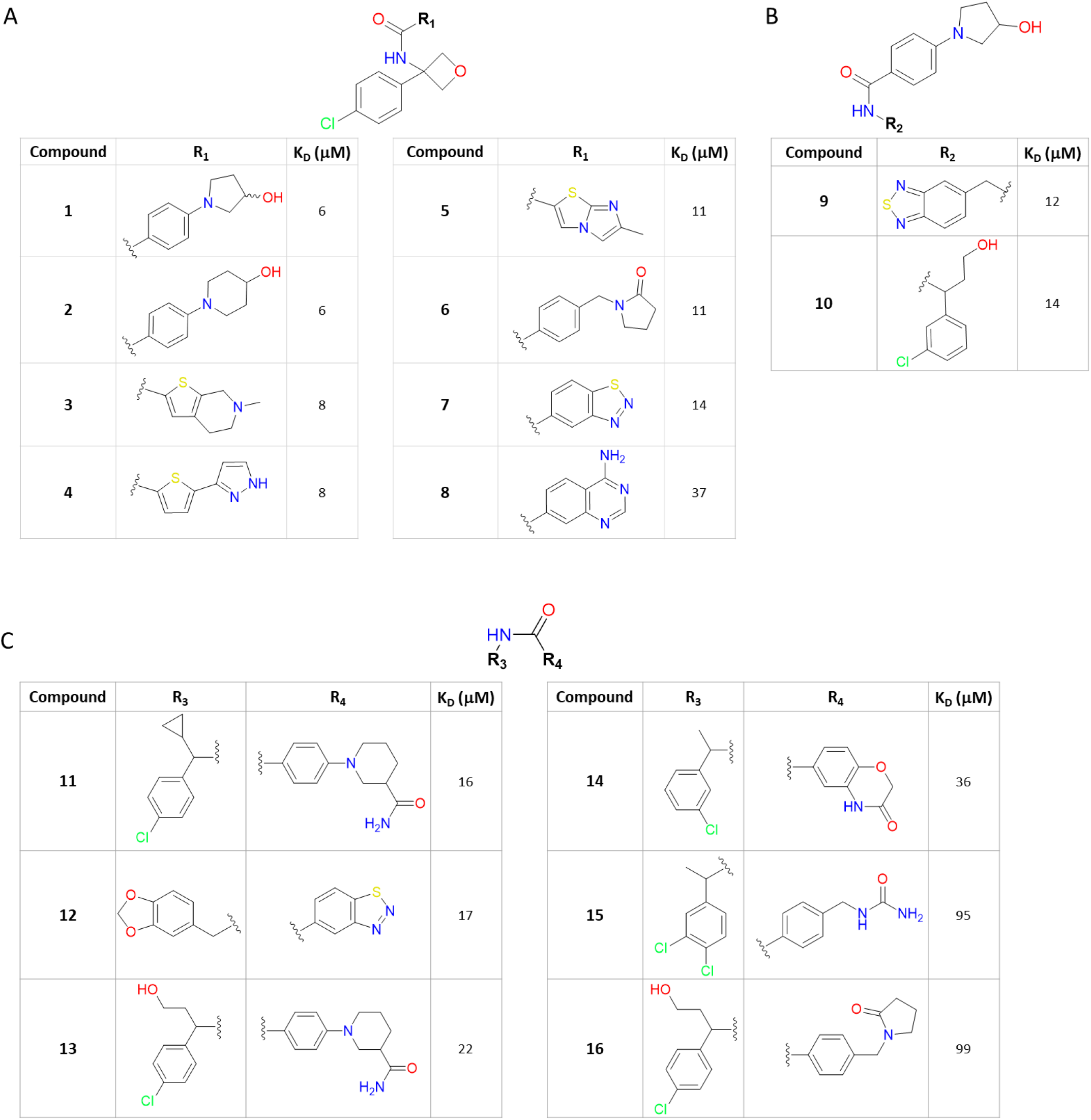
Machine-learning-assisted SAR-by-catalog. Sixteen analogs which explore the tolerance of various functional groups on (**A**) right-hand side, (**B**) left-hand side, and (**C**) both right- and left-hand side of the amide bond.

**Figure 3:**
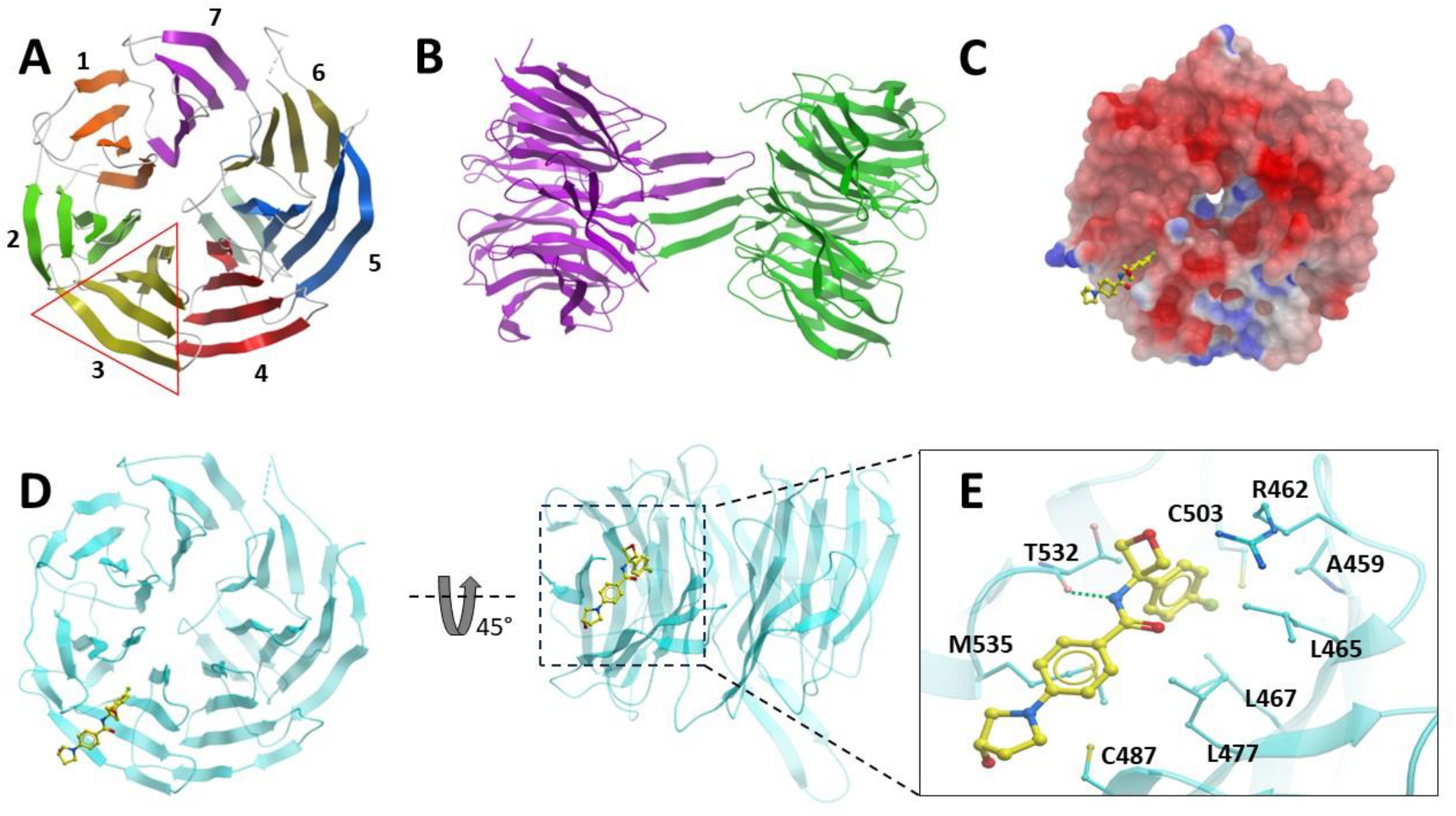
Crystal structures of WDR91 apo and in complex with compound 1. (**A and B**) Cartoon representation of apoWDR91 (PDB ID: 6VYC), forming homodimers in the crystal asymmetric unit (shown in B). (**C**) Electrostatic surface potential representation of WDR91 in complex with compound **1**. WDR91 surface color indicates electrostatic potential ranging from −5 kT e^−1^ (red) to +5 kT e^−1^ (blue). Electrostatic surface potentials were calculated in ICM. (**D**) Cartoon representation of WDR91 bound to **1** (yellow sticks), revealing the compound binding side pocket of WDR domain. (**E**) Close-up view of the WDR91 side pocket highlighting the key residues interacting with compound **1**. The hydrogen-bonding interaction between the backbone oxygen of threonine (T532) and the amide nitrogen of the compound is shown with a dotted green line.

### Crystal structures of apo WDR91 and in complex with compound **1**

To further elucidate the interaction of compound **1** with WDR91, we determined the crystal structures of the WDR domain of WDR91 in apo form (containing a loop deletion at residues 522-536), and in complex with **1** at 2.1 and 2.2 μ resolution, respectively. These structures are hereafter referred to as apoWDR91 and WDR91-**1** (PDB IDs: 6VYC and 8SHJ). Table 1 summarizes the crystallographic data collection, refinement, and validation statistics. The fifteen-residue loop deletion within the WDR domain of WDR91 was designed to improve the crystal packing and to obtain diffraction-quality crystals, which wasn’t amenable with the WT protein. In apoWDR91, the WDR domain adopts a seven-bladed β-propeller fold, as commonly observed in other WDR proteins **(Fig. 3A)**^17,26^. In apoWDR91, the crystal asymmetric unit contains two WDR91 molecules forming a head-to-head dimer mediated by a β-hairpin loop (residue range 652-667) located at the top side of the WDR domain **(Fig. 3 B)**. Whereas in WDR91-**1**, there are three WDR91 molecules in the crystal asymmetric unit, in which, two of them form a head-to-head dimer, similarly to the dimer observed in apoWDR91, but the third molecule remains unpaired. The latter has a higher overall B-factor value compared to the other two molecules, presumably due to the absence of a stabilization effect by dimerization. Well-resolved electron density map was observed for compound **1** in all three molecules of WDR91-**1**, revealing compound binding to a side pocket located at the bottom side of WDR91 (**Fig. 3C, D**). Several leucine (L465, L467 and L477), an alanine (A459) and methionine (M535) residues were found to be involved in making hydrophobic interactions with the compound. The 4-chlorophenyl moiety of **1** binds deep inside the groove where it forms a halogen bond with the thiol group of C503.

**Table 1.** Data collection and refinement statistics.

|  | apoWDR91 | WDR91-1 | WDR91-18 |
| --- | --- | --- | --- |
| PDB ID | 6VYC | 8SHJ | 8T55 |
| Wavelength (Å ) | 1.5418 | 0.97918 | 0.97918 |
| Resolution range (Å) | 46.07-2.10 (2.16-2.10) * | 46.25-2.21 (2.25-2.21) * | 38.78-2.20(2.24-2.20) * |
| Space group | P2<sub>1</sub>22<sub>1</sub> | P2<sub>1</sub>2<sub>1</sub>2<sub>1</sub> | P2<sub>1</sub>2<sub>1</sub>2<sub>1</sub> |
| Unit cell (Å) | 77.0, 84.0, 110.2 | 76.9, 121.4, 131.8 | 76.7, 121.6, 132.0 |
| Total reflections | 332722 | 463068 | 478836 |
| Unique reflections | 41340 (3269) | 62725 (3080) | 62021 (3075) |
| Multiplicity | 8.0 (8.3) | 7.4 (7.3) | 7.7 (8.2) |
| Completeness (%) | 97.6 (95.6) | 99.7 (99.5) | 97.9 (98.4) |
| Mean I/sigma(I) | 16.8 (2.8) | 17.1 (1.5) | 23.0 (1.9) |
| Wilson B-factor | 39.5 | 36.2 | 46.2 |
| R-merge | 0.091 (0.906) | 0.099 (0.992) | 0.080 (1.014) |
| R-meas | 0.098 (0.966) | 0.106 (1.068) | 0.086 (1.083) |
| R-pim | 0.034 (0.331) | 0.039 (0.391) | 0.032 (0.375) |
| CC1/2 | 0.999 (0.827) | 0.996 (0.744) | 0.998 (0.826) |
| Reflections used in refinement | 39208 | 62656 | 60770 |
| Reflections used for R-free | 2085 | 1224 | 1207 |
| R-work | 0.221 | 0.211 | 0.245 |
| R-free | 0.251 | 0.257 | 0.288 |
| Number of non-hydrogen atoms |  |  |  |
| Macromolecules | 4906 | 7232 | 7212 |
| Ligands | n/a | 78 | 85 |
| Solvent | 127 | 265 | 235 |
| Protein residues |  |  |  |
| RMS (bonds) | 0.005 | 0.010 | 0.007 |
| RMS (angles) | 1.348 | 1.13 | 1.270 |
| Ramachandran favoured (%) | 97.4 | 96.8 | 96.4 |
| Ramachandran allowed (%) | 99.7 | 100 | 100.0 |
| Ramachandran outliers (%) | 0.2 | 0 | 0 |
| Rotamer outliers (%) | 2.52 | 4.98 | 1.79 |
| Clash score |  |  |  |
| Average B-factor |  |  |  |
| Macromolecules | 44.5 | 46.6 | 64.6 |
| Ligands | n/a | 50.1 | 77.5 |
| Solvent | 37.3 | 265 | 50.1 |

\*Statistics for the highest-resolution shell are shown in parentheses.

A neighboring arginine side chain (R462) forms a hydrogen-bond with the oxetane oxygen of **1** in chain B, but is not observed in other chains, suggesting that this interaction is not crucial for compound binding. The backbone oxygen atom of T532 forms a hydrogen bonding interaction with the amide nitrogen of **1** (**Fig. 3E**), thus shifting the 532-534 loop region ∼2.2μ closer to the amide nitrogen of **1** compared to apoWDR91 (**Fig. S1A**).

### Identification and orthogonal confirmation of covalent ligands

In WDR91-**1**, the pyrrolidine ring moiety of **1** is located near cysteine 487 at the entrance of the compound binding pocket, which could be exploited to convert the compound into a covalent analogue (**Table S2** entries 17-20). However, no close analogs of **1** with a covalent warhead at the pyrrolidine ring region were commercially available, except for compounds **17-20** with acrylamide warhead, which share some core structural features with **1**. In **17** and **20**, the *p*-chlorophenyl group is replaced with phenyl and 4-fluorophenyl groups, respectively. Compounds **18-20** feature tetrahydroquinoline amides instead of phenyl amides. In addition, compounds **19** and **20** have cyclopropyl and cyclohexyl groups, respectively, instead of oxetane.

Compounds **17-20** were tested for covalent adduct formation using intact mass LC-MS. WDR12 was tested in parallel with ctdWDR91 to evaluate selectivity. LC-MS was performed after incubating the protein with each of the compounds **17-20** for 2 hours. Compounds **18** and **19** showed adduct formation with ctdWDR91 (**Fig. 4**), but not with the negative control protein WDR12 (**Fig. S2A-B**). Compounds **17** and **20** did not appear to form a covalent adduct with either protein.

**Figure 4:**
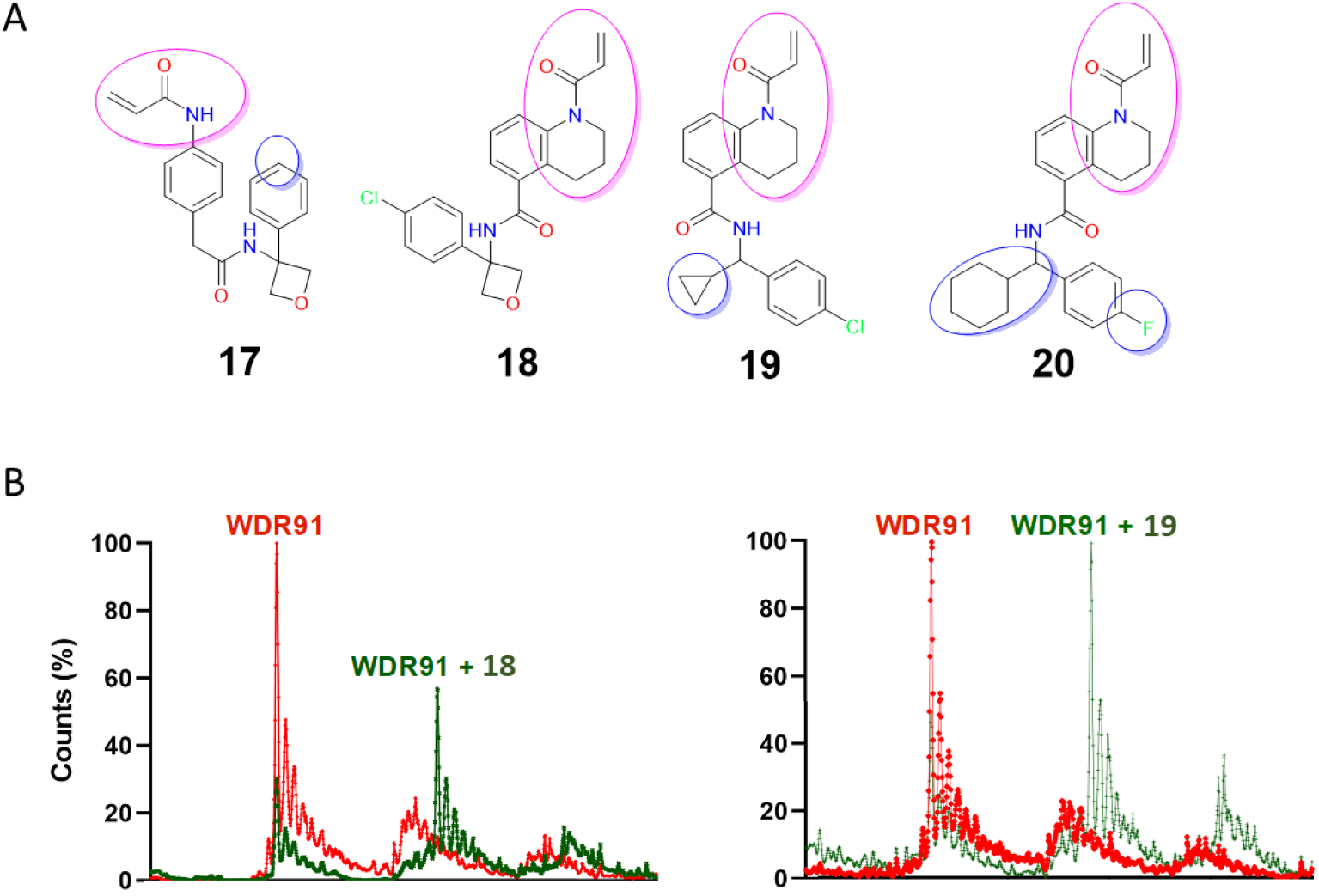
Covalent analogs and screening by intact mass LC-MS. (**A**) Chemical structures of covalent analogs (**17-20**) of the co-crystallized hit **1**, and (**B**) intact mass LC-MS of ctdWDR91 in complex with **18, 19** (molar ratio of protein to ligand is 1:10) where the two major peaks correspond to the apo (red) and the complex (green), respectively.

Thermal stability and SPR were used for orthogonal validation of the covalent analogs **18** and **19**. The ctdWDR91 was incubated for 2 hours in the absence and presence of compounds **18** and **19** (with compound concentrations ranging from 0 to 200 mM). Both compounds displayed sigmoidal dose-response curves with ∼*T*_m_ ∼4 °C at saturation (**Fig. 5A-C**), and in a dose-dependent manner. Compounds **18** and **19** were also tested by SPR and yielded *K*_D_ values 43 ± 7 μM and 47 ± 8 μM, respectively (**Fig. 5D-F**). The dissociation of compounds **18** and **19** during SPR can be explained by the fast compound contact time (60s), which presumably is not enough for covalent adduct formation with C487, suggesting that covalent adduct formation is a secondary event for compounds **18** and **19** following non-covalent binding to WDR91. On the other hand, compounds **17** and **20** did not show any significant binding to ctdWDR91 by SPR (**Fig. S4A-C**).

**Figure 5:**
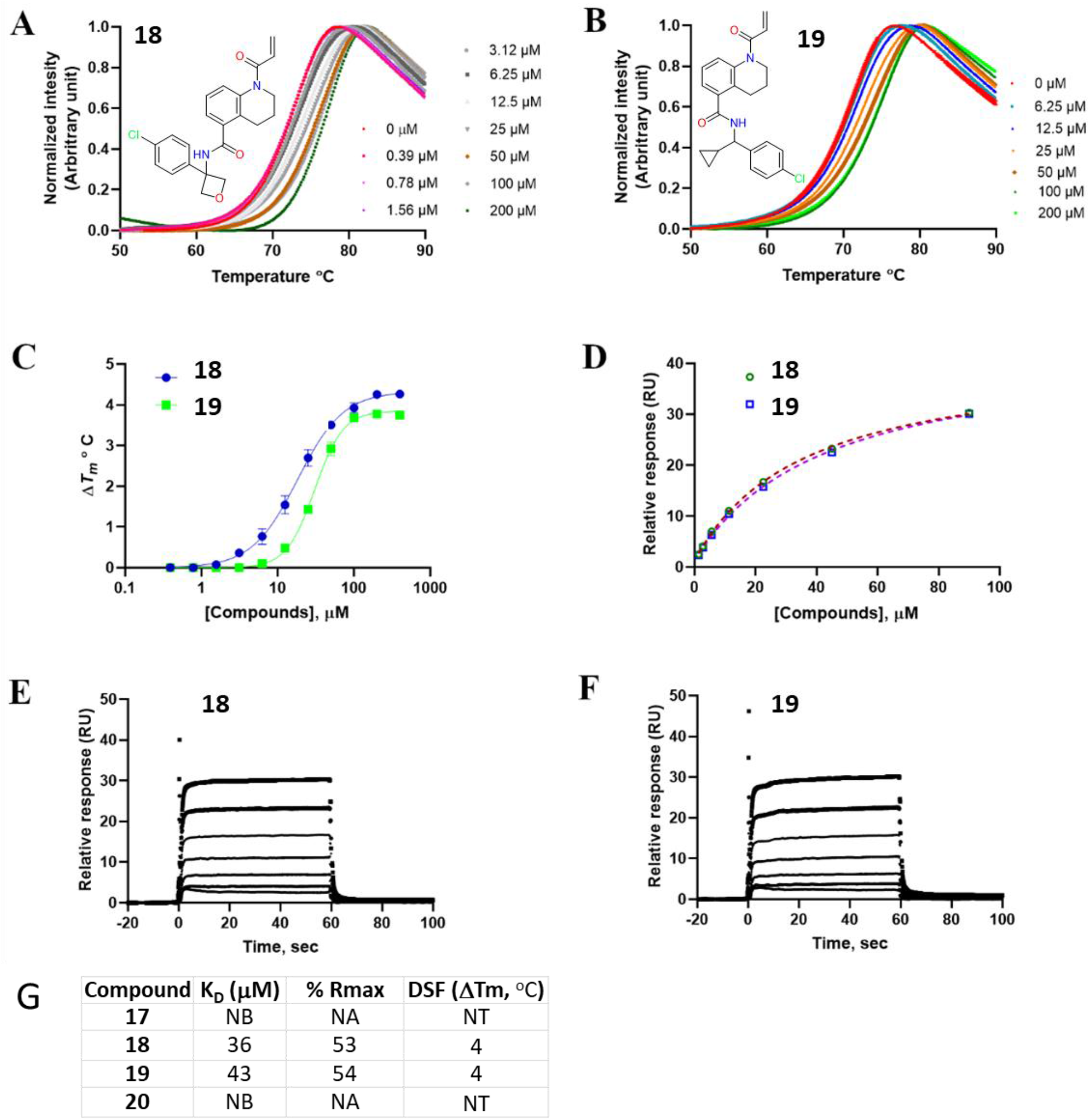
Thermal stabilization of WDR91 in the presence and absence of covalent hits at varying compound concentrations. (A, B and C) The compounds showed thermal stabilization of WDR91 complexes upon binding to ctdWDR91 in a dose-dependent manner. **Affinity determination by SPR**. (**D**) Relative response vs concentration plots for affinity determination, (**E, F**) Raw sensorgrams showing binding of both covalent compounds to ctdWDR91. (**G**) A summary of the primary screening and orthogonal confirmation of binding. NB = no binding, NA = not applicable, NT = not tested (Data from one replicate is shown here for clarity).

### Co-crystal structure of WDR91 with a covalent analog

The co-crystal structure of WDR91 in complex with **18** (hereafter referred to as WDR91-**18**) was determined at 2.2Å resolution in P2_1_2_1_2_1_ space group with three WDR91 molecules in the crystal asymmetric unit (PDB ID: **8T55)**. Similar to WDR91-**1**, the two WDR91 molecules in WDR91-**18** (chains A and B) form head-to-head dimers with each other, whereas the third molecule (chain C) does not form any dimeric interaction and has a poor overall electron density map, which was not reliably built, thus likely contributes to the overall high Rfactor/Rfree values observed for WDR91-**18**. Accordingly, the electron density map for compound **18** was best observed in chains A and B **(Fig. S3)**, which are used here for structural characterization. The covalent compound **18** occupies the same pocket as **1**, and the residues lining the binding pocket align well in both structures (**Fig. 6A, B**). As expected, the acrylamide warhead of **18** forms a covalent adduct with the thiol group of C487 (*via* a Michael addition) within the same binding pocket (**Fig. 6C**). The 4-chlorophenyl, oxetane ring and the central amide are in the same position as observed with **1**. Indeed, the complex retains some of the key interactions e.g., the halogen bond between the chlorine (acceptor) and the sulfur (donor) in C503, and the hydrogen-bond between amide nitrogen in **18** and the backbone amide oxygen of T532.

**Figure 6:**
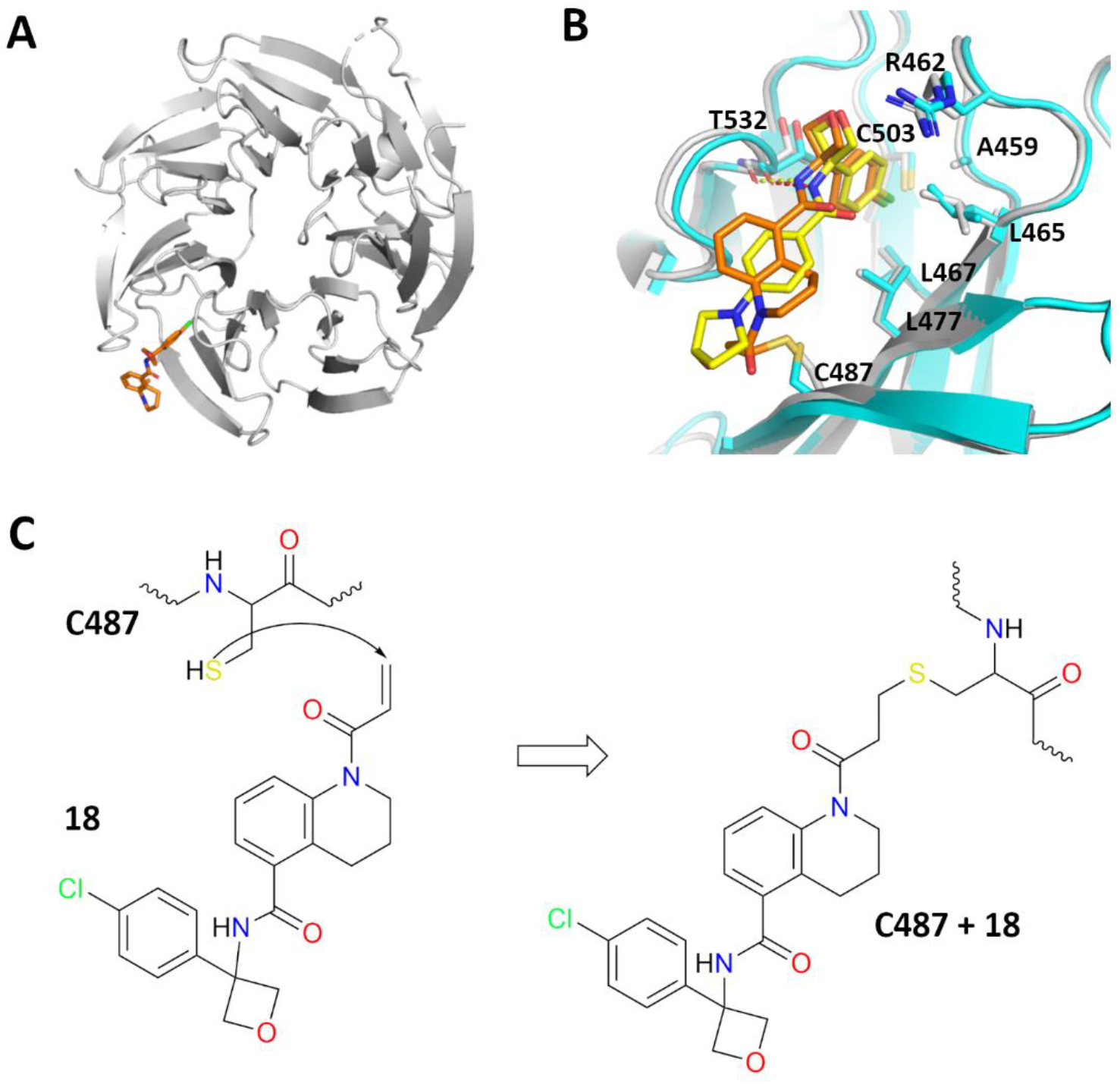
Co-crystal structure of WDR91 in complex with covalent compound 18. (**A**) Cartoon representation of WDR91-**18** complex showing the compound (orange) binding pocket. (**B**) Superposition of two WDR91 co-crystal structures, one in complex with **1** (yellow) and the other with the covalent compound **18** (Orange). A zoomed-in view of the compound binding pocket showing the bound ligands, **1** (yellow) and **18** (orange) and the residues involved in the interaction are shown in cyan and grey, respectively. (**C**) Schematic of covalent adduct formation between the thiol group of C487 and the acrylamide warhead of **18**.

These biophysical and structural data describe novel small molecule side-pocket binders of the WDR domain of WDR91. WDR domains are protein interaction hubs and are known to interact with their respective substrates using the central and side pockets ^17^. It has already been shown that the WDR domain-containing protein WDR5 uses its central channel to interact with the histone methyltransferase MLL1, whereas the side pocket is used to recruit an essential component of the MLL1 complex, RBBP5^27^. A long-range allosteric effect has also been reported in the case of yeast Cdc4, where the binding of a small molecule ligand in a side pocket of WDR domain modulates the interactions in the central channel^28^. Previous reports indicate that WDR91 uses the bottom side of the WDR domain to interact with Rab7a^6^. However, the physiological implications of the compound-binding side pocket of WDR91 are yet to be explored.

## Conclusions

We report here the discovery of first-in-class WDR91 small molecule ligands using affinity-mediated DNA-encoded chemical library (DEL) selection followed by model-building using machine-learning. This model was used for predicting primary hits within a virtual catalog, followed by SAR-by-catalog. We demonstrate that training of an ML model using DEL selection output data can enable and accelerate the discovery of small molecule ligands from readily accessible compounds, such as the Enamine REAL database, without the need for expensive, custom, off-DNA chemical synthesis. Through this approach, we identified the hit compound **1**, which was validated by SPR to have a K_D_ value of 6 ± 2 μM. We further characterized the binding mode of **1** by X-ray crystallography, revealing that the compound binds to a side pocket of WDR91 nestled between the β-propeller blades 3 and 4. Structure-guided optimization of **1** binding to WDR91 facilitated the design of covalent analogs bearing a reactive acrylamide warhead at a position adjacent to cysteine 487 in the compound binding pocket of WDR91, which supported covalent adduct formation. The covalent binding modes for both compounds **18** and **19** were confirmed by intact mass LC-MS, and further characterized by SPR, DSF and X-ray crystallography. These reversible and irreversible chemical compounds reported here could guide medicinal chemistry efforts to develop potent and selective WDR91 chemical probes to further characterize its function in cellular disease models and evaluate its therapeutic potential as an antiviral against coronaviruses and related viruses.

## Abbreviations

TPU: tensor processing unit
GCNN: graph convolution neural network
SAR: structure activity relationship
DEL: DNA-Encoded Chemical Library
ML: machine learning
DEL-ML: DNA Encoded Chemical Library selection followed by machine learning
LC-MS: liquid chromatography mass spectrometry.

## Experimental Section

### Affinity-Mediated Selection of the DNA-Encoded Chemical Library Deck

The purified WDR91 protein (construct JMC067-F09) containing an N-terminal His_6_-tag at a concentration of 4.1 uM was incubated with the DEL deck as described in Cuozzo et al 2017^24^ at a total concentration of 40 μM for 1 h in a volume of 60 μl in 1× selection buffer (8 mM NaOAc, 134 mM KOAc, 4 mM NaCl, 0.8 mM Mg(OAc)2, 5 mM imidazole, 1 mM TCEP, 1 mg/mL sheared salmon sperm DNA, 0.02% Tween20, 20 mM HEPES, pH7.2). A selection condition containing no target was performed in parallel to the selection condition containing target protein utilizing all the same conditions (except for presence of input protein) to evaluate matrix binders. For each separate selection, a separate ME200 tip (Phynexus) containing IMAC affinity matrix was prewashed three times in 200 μL of fresh 1× selection buffer. Each incubated sample was separately captured with 20 passages over the ME200 tip over a duration of 0.5 h. The bound protein and associated library members were captured on the ME200 tip and then washed eight times with 200 μL of fresh 1× selection buffer. Bound library members were eluted by incubating the ME200 tip with 60 μL of 1× fresh selection buffer at 85°C for 5 min. The eluted sample was then passed over a fresh, prewashed ME200 tip containing appropriate affinity matrix to remove any eluted protein over 20 passages. A second round of selection was conducted using the eluate of the first selection and appropriate protein reagents. The eluate of the second round of selection was PCR amplified and sequenced as described in Cuozzo et al 2017^24^. Sequence data were parsed, error-containing sequences were disregarded, amplification duplicates were removed, and building block combinations and chemical scheme encodings were decoded and reported along with associated calculated statistical parameters.

Capture of proteins during selection was determined by analyzing selection samples by SDS-PAGE. Nine μL of input and resin samples were mixed with 3 μL of 4x denaturing load buffer, heated at 95°C for 5 min then run on a 4-12% Bis-Tris gel (BioRad 3450125). The input sample is identical to the initial solution containing protein in 60 μL 1x selection buffer prior to capture with 20 passages over the ME200 tip. The resin sample is the resin removed from the ME200 tip and mixed to resuspension in 60 μL 1x selection buffer and gel-loading buffer. SDS-PAGE data are shown in Figure S5.

## Machine learning

### Label assignment

Sequence data were parsed, error-containing sequences were discarded, amplification duplicates were removed, and building block and chemical scheme encodings were decoded and reported along with associated calculated statistical parameters. These chemical data were aggregated into disynthons and enrichment values were calculated as described in McCloskey et al^23^. Disynthons were classified into four labels: TARGET_HIT, MATRIX_BINDER, PROMISCUOUS_BINDER, and NON-HIT. The label derivation is a multi-class decision tree, as shown in Figure 1 of Han et al.^29^. The positive class for machine learning is TARGET_HIT, which included disynthons enriched in the target condition but were not enriched in the matrix binder condition. Disynthons that have a history of enrichment across targets were labeled as promiscuous binders and removed from the TARGET_HIT class. In total, our model training comprised approximately 250 million labeled examples.

### Model training

The Graph Convolutional Neural Network (GCNN) model in this study is similar to the one described in McCloskey et al.^23^, with the following improvements. The model training process described in McCloskey et al.^23^ and Kearnes et al.^30^ has been migrated to tensor processing units (TPU) to take advantage of TPU’s strength in large batch synchronous training. The use of TPUs resulted in a significant reduction in time to training convergence. The training time was reduced from more than 1 week to about 30 hours. This enabled the authors to perform 5-6 times more experiments. Most hyperparameters of our model are the same as the “W2N2” variant specified in Kearnes et al.^30^, with the following updates: 1) learning rate was set to 0.1, 2) training step was set to 1,000,000 steps with a checkpoint saved every 100,000 steps, 3) batch size was 168, 4) training data was split into 23 folds.

### Model Ensembling

In addition to ensembling the cross-validation fold models as described in McCloskey et al.^23^, we introduced two additional levels of model ensembling to improve the overall reliability of our model. To reduce the variance in the final prediction, we used TPU supported graph partitioning to do manual model parallelism. We trained 8 models with independently randomly initialized sets of model weights, each trained on its own TPU core. The final prediction is the median of the predictions from the 8 models. We further reduced the variance in the final prediction by repeating the entire training process 3 times and then by ensembling the replicas. The final prediction score is the median of the predictions from the 3 training runs, each with 8 TPU model replicas for every one of the 23 cross-validation folds.

### Model step selection

Unlike the maximum ROC-AUC metric described in McCloskey et al.^23^, we used the top_100_actives metric to perform model selection. The top_100_actives metric is calculated by counting how many active examples are in the top 100 compounds ranked by their hit class prediction value. After training a model for each cross-validation fold split, the model weights of the saved checkpoint with the maximum top_100_actives for the hits class on the tuning set were selected. In other words, we selected the model checkpoint for each cross-validation fold split that produced the highest number of hit compounds in the top 100 tuning set predictions.

### Property filtering and Diversity selection

Property filtering and diversity selection were applied to the top model predictions, resulting in 200 diverse compounds for purchasing. Specifically, we used the rd_filters open-source package (https://github.com/PatWalters/rd_filters) to remove molecules that contain undesirable substructures. In addition, we applied molecular weight (MW) > 250 and a Solubility Forecast Index (SFI) < 6.5 filters. Lastly, a directed sphere exclusion (DISE) algorithm^31^ selected the highest scoring molecules of each cluster for purchasing. For DISE, we used Morgan Fingerprint with radius of 3 (ECFP6) and a Tanimoto similarity cutoff of 0.3. To ensure scaffold diversity, each selected compound is a unique Murcko scaffold.

One hundred compounds were selected from Mcule Instock (9.35M compounds) and 100 compounds were selected from Enamine REAL (1.9B compounds). Mcule and Enamine successfully synthesized 72 and 78 compounds, respectively.

### DEL-ML-driven hit expansion

In this study, we conducted an automated hit-expansion campaign utilizing GCNN model-guided search and prioritization to identify analogs of compound **1**. The search was performed against a 1.9 billion commercially available Enamine REAL library that did not require expensive and time-consuming bespoke synthesis. First, we selected 50 diverse analogs with an ECFP6 Tanimoto similarity greater than 0.55. This process was sampled near neighbors of compound **1**. Next, we relaxed the Tanimoto similarity criteria to 0.4 and selected an additional 50 diverse analogs. This process allowed the GCNN model to explore chemical space that is of moderate distance from compound **1**. A DISE radius of 0.2 was used for diversity selection. Of the 100 selected molecules, duplicates were removed resulting in 86 ordered compounds. Enamine successfully synthesized and delivered 84 molecules.

### Protein expression and purification

#### Biotinylated ctdWDR91 used in SPR

DNA fragments encoding full-length WDR91 (M1-A747) and truncated WDR91 (P392-A747) were PCR amplified from cDNA (MGC:29694) and cloned into the pFBD-BirA expression vector. pFBD-BirA is a derivative of pFastBac Dual (Invitrogen) that co-expresses biotin ligase and adds an N-terminal biotin acceptor peptide and a C-terminal 6X His-Tag to WDR91. The resulting plasmid was transformed into DH10Bac™ Competent E. coli (Invitrogen) and a recombinant viral DNA bacmid was purified and followed by a recombinant baculovirus generation in sf9 insect cells^32^. Scale-up production of the biotinylated ctdWDR91 has been done with addition into baculovirus infected Sf9 cells culture of D-BIOTIN (Sigma-Aldrich) to the final concentration of 10 μg/mL. The cells were collected after 72-96 hours post infection with well-developed signs of infections and 70-80 % viability. The harvested cells were re-suspended in binding buffer containing 20mM Tris-HCl, pH 8.0 containing 500 mM NaCl, 5mM imidazole and 5% glycerol, 0.5 mM TCEP, 1X protease inhibitor cocktail (100 X protease inhibitor stock in 70% ethanol (0.25 mg/mL Aprotinin, 0.25 mg/mL Leupeptin, 0.25 mg/mL Pepstatin A and 0.25 mg/mL E-64) or Roche complete EDTA-free protease inhibitor cocktail tablet. The cells were lysed chemically by rotating for 30 min with NP40 (final concentration of 0.5%) and 22.5 U/mL Benzonase nuclease (in house) followed by sonication at frequency of 7.5 (10” on/7” off) for 5 min (Sonicator 3000, Misoni). The crude extract was clarified by high-speed centrifugation (60 min at 28,000 g at 4°C) by Beckman Coulter centrifuge. The cleared lysate was loaded onto Ni-NTA affinity resin column (Qiagen) pre-equilibrated with binding buffer. The column was washed with binding buffer containing 30 mM imidazole after washing the column with 1 mM biotin in PBS. Subsequently, the column was eluted in binding buffer containing 250 mM imidazole. The purity of the fractions was confirmed on SDS-PAGE gels (NuPAGE 4-12% Bis-Tris gel; Invitrogen) before loading the protein onto a Superdex200 26/600 size-exclusion column for further purification.

#### Protein used in Crystallography and Assays

DNA fragments encoding the WDR91 WDR domain (P392-A747) and the WDR91 WDR domain with a loop deleted (P392-A747 with residues Q522-N536 removed) were PCR amplified from cDNA (MGC:29694). Each fragment was cloned into the pFBOH-MHL expression vector (Addgene Plasmid #162266) and are referred to here as WDR91 (WD) and xtalWDR91 respectively. The baculovirus expression vector system (BEVS) have been used to express these recombinant proteins. The harvested cells were lysed by sonication in a buffer containing 20 mM Hepes pH 7.4, 500 mM NaCl, 5% glycerol, 0.06% NP40, Benzonase, protease inhibitor cocktail. The protein was initially affinity purified by TALON Metal Affinity Resin (Cat# 635504 Clontech). The eluted fractions were then further purified by size-exclusion chromatography on a Superdex200 26/60 column and eluted with a buffer containing 20 mM Tris-HCl pH 8.0 and 150 mM NaCl. The protein purity and quality were assessed by SDS-PAGE and Mass spectrometry. The pure fractions were pooled together and concentrated to 8.44 mg/mL, flash frozen in liquid nitrogen and stored in -80 °C freezer for future crystallization trials.

### Surface plasmon resonance (SPR)

SPR screening were performed using both Biacore T200 and Biacore 8K instruments at 20 °C. Biotinylated ctdWDR91 (392-747) was immobilized on a SA sensor chip using two separate active flow cells, while the remaining active flow cell was charged with a negative control protein, WDR5, after conditioning both reference and active flow cells with 50 mM NaOH for 3 X 60 s with a flow rate of 10 μl/min. During the protein immobilization, each protein solution (30 μg/mL) was injected through the respective active flow cells for 60s with a flow rate of 5 μl/min, which resulted in an immobilization level of 6000-7500 RU. After protein immobilization in the active flow cells, equilibration was performed by flowing running buffer (10 mM HEPES, pH 7.4, 150 mM NaCl, 3 mM EDTA, 0.03% Tween 20 and 3% DMSO) over the flow cells with a flow rate of 50 μl/min until a stable base line was observed. The compounds were then injected over the reference and active flow cells using multi-cycle kinetics at a flow rate of 40 μL/min with a 60s association and 120s dissociation time for single concentration screening. On the other hand, single-cycle kinetics with a five successive injection with a 60 s association at a flow rate of 40 μL/min and a 120s dissociation after the final injection was performed for the dose-response experiments. To assess the covalent compounds, dose-response titrations were performed using both Biacore T200 and Biacore 8K instruments where biotinylated ctdWDR91 (392-747) were immobilized with an RU of ∼ 4000-6000 on a Neutravidin coupled CM5 sensor chip. The covalent compounds and compound **1** (as a control) were injected over the reference and active flow cells using multi-cycle kinetics at a flow rate of 45 μL/min with a 60 s association and 60 s dissociation time. For both multi and single cycle kinetics, 10 start-ups, blanks and wash (50% DMSO wash to flush the needles) cycles were included before and after each cycle. The compound dilution was performed using the running buffer by maintaining the final DMSO concentration of 3% (v/v) across the tested concentration range. Eight-point solvent correction was also included for each run to adjust high bulk responses from the solvent. Double referencing of the data was performed by subtraction of the reference flow cell and then the respective blank cycles. The data analysis was performed using BIA evaluation software and affinity fitting was performed by applying 1:1 binding model provided with the software. The final figures were prepared by exporting the data to GraphPad Prism.

### Differential scanning fluorimetry (DSF)

The thermal stability of WDR91(WD) proteins in the presence/absence of hit compounds were assessed by differential scanning fluorimetry. The experiments were performed in 384-well PCR plate format using a reaction mixture of 20 μL. The protein samples, 0.1 mg/ml of WDR91(WD) and/or WDR91 (WD loop deleted), were mixed with/without compounds at varying compound concentrations ranging from 0-200 μM in the assay buffer (100 mM HEPES, pH 7.5, and 150 mM NaCl) containing SYPRO Orange (1:1000) with a final DMSO concentration of 2% (v/v). A two-hour pre-incubation step was also introduced in the case of covalent compounds. After a brief centrifugation, the reaction mixture was subjected to temperature scan between 25 °C to 95 °C with a 4°C increment/min using a real-time PCR instrument (LightCycler 480 from Roche). The data were extracted and analyzed by fitting the thermal denaturation curve to the Boltzmann equation to get the *T*_m_ (melting temperature) by non-linear regression using BioActive 2.1.10.

### Mass spectrometry

The protein samples were pre-mixed with/without compounds in 20 mM HEPES, pH 7.5, 150 mM NaCl and incubated for 2 hours at room temperature prior analyzing on LC/MS to identify potential covalent ligands. Total 10 μM of WDR91 (WD) or WDR12 (WD) proteins were used in the assay mixture with variable concentrations of compounds by maintaining 1:1, 1:5, and 1: 10 proteins to compound ratio. The assay mixture for each sample were quenched by 1% trifluoroacetic acid after 2 hrs of incubation and separated over an Agilent 1260 capillary HPLC system before acquiring MS data using an Agilent Q-TOF 6545 mass spectrometer via dual Agilent Jetstream ion source. The raw data were analyzed by Agilent MassHunter Qualitative Analysis B.07.00.

### Crystallization, Data collection, Processing and Refinement

The xtalWDR91 protein was used for generating both apoWDR91 structure and in complex with small molecule modulators. The co-crystallization was performed using the loop-deleted truncated version of WDR91 and hit compounds. Initially, the crystallization screening was carried out in 96-well plate format by sitting drop vapor diffusion method using two in-house screens and the plates were set up by a liquid handler with a protein to reservoir solution ratio of 1:1. Based on the initial crystal hits, grid optimization was performed in 24-well plate format by varying different components of the selected conditions, e.g., pH, salt and precipitant concentrations, by hanging drop vapor diffusion method. Crystals were grown at room temperature for several weeks. For all crystallization experiments, 0.25 mM and/or 1.2 mM of compounds were added to 6 mg/ml of protein solution with a final DMSO concentration of 2-5% (v/v) and centrifuged at 14000xg for 10 minutes at 4 °C prior to setting up the drops. The matured crystals were harvested after 1-2 weeks using the respective reservoir solution containing 10% glycerol as a cryo-protectant. Table 2 summarizes the crystallization conditions, data processing and refinement information for all three WDR91 structures reported in this manuscript.

**Table 2.** Summary of WDR91 crystallization condition.

|  | apoWDR91<br>(PDB ID: 6VYC) | WDR91-1<br>(PDB ID: 8SHJ) | WDR91-18<br>(PDB ID: 8T55) |
| --- | --- | --- | --- |
| Crystallization condition | 12% PEG20000, 0.1 M MES, pH 6.5 | 12% P3350, 0.2 M KCl, 0.1 M MES, pH 6.5 | 12% P3350, 0.2M KCl, 0.1 M MES, pH 6.5 |
| Protein: compound ratio | N/A | 1:2 | 1:10 |
| Cryo condition | 10 % glycerol | 10 % glycerol | 10 % glycerol |
| Temperature, ° K | 293 | 293 | 293 |

## Acknowledgement

We would like to thank Dr. Fengling Li for her assistance with the biophysical experiments, and Maria Kutera and Zahra Hejazi from SGC-Toronto for protein expression in Sf9 insect cells. We thank the staff at the Northeastern Collaborative Access Team, which is funded by the National Institute of General Medical Sciences from the National Institutes of Health (P30 GM124165). The Eiger 16M detector on the 24-ID-E beamline is funded by an NIH-ORIP HEI grant (S10OD021527). This research used resources of the Advanced Photon Source, a U.S. Department of Energy (DOE) Office of Science user facility operated for the DOE Office of Science by Argonne National Laboratory under Contract No. DE-AC02-06CH11357. The Structural Genomics Consortium is a registered charity (no: 1097737) that receives funds from Bayer AG, Boehringer Ingelheim, Bristol Myers Squibb, Genentech, Genome Canada through Ontario Genomics Institute [OGI-196], EU/EFPIA/OICR/McGill/KTH/Diamond Innovative Medicines Initiative 2 Joint Undertaking [EUbOPEN grant 875510], Janssen, Merck KGaA (aka EMD in Canada and US), Pfizer and Takeda.

## Accession Numbers

Atomic coordinates and structure factors for the reported crystal structures have been deposited in the Protein Data bank under the accession numbers: **6VYC, 8SHJ, 8T55**.

## References

(1) Langemeyer, L.; Fröhlich, F.; Ungermann, C. Rab GTPase Function in Endosome and Lysosome Biogenesis. Trends Cell Biol 2018, 28 (11), 957–970. https://doi.org/10.1016/j.tcb.2018.06.007.

(2) Liu, K.; Jian, Y.; Sun, X.; Yang, C.; Gao, Z.; Zhang, Z.; Liu, X.; Li, Y.; Xu, J.; Jing, Y.; Mitani, S.; He, S.; Yang, C. Negative Regulation of Phosphatidylinositol 3-Phosphate Levels in Early-to-Late Endosome Conversion. J Cell Biol 2016, 212 (2), 181–198. https://doi.org/10.1083/jcb.201506081.

(3) Casanova, J. E.; Winckler, B. A New Rab7 Effector Controls Phosphoinositide Conversion in Endosome Maturation. J Cell Biol 2017, 216 (10), 2995–2997. https://doi.org/10.1083/jcb.201709034.

(4) Christoforidis, S.; Miaczynska, M.; Ashman, K.; Wilm, M.; Zhao, L.; Yip, S. C.; Waterfield, M. D.; Backer, J. M.; Zerial, M. Phosphatidylinositol-3-OH Kinases Are Rab5 Effectors. Nat Cell Biol 1999, 1 (4), 249–252. https://doi.org/10.1038/12075.

(5) Robinson, F. L.; Dixon, J. E. Myotubularin Phosphatases: Policing 3-Phosphoinositides. Trends Cell Biol 2006, 16 (8), 403–412. https://doi.org/10.1016/j.tcb.2006.06.001.

(6) Liu, K.; Xing, R.; Jian, Y.; Gao, Z.; Ma, X.; Sun, X.; Li, Y.; Xu, M.; Wang, X.; Jing, Y.; Guo, W.; Yang, C. WDR91 Is a Rab7 Effector Required for Neuronal Development. Journal of Cell Biology 2017, 216 (10), 3307–3321. https://doi.org/10.1083/jcb.201705151.

(7) van der Beek, J.; de Heus, C.; Liv, N.; Klumperman, J. Quantitative Correlative Microscopy Reveals the Ultrastructural Distribution of Endogenous Endosomal Proteins. J Cell Biol 2022, 221 (1). https://doi.org/10.1083/jcb.202106044.

(8) Bouhaddou, M.; Memon, D.; Meyer, B.; White, K. M.; Rezelj, V. V; Correa Marrero, M.; Polacco, B. J.; Melnyk, J. E.; Ulferts, S.; Kaake, R. M.; Batra, J.; Richards, A. L.; Stevenson, E.; Gordon, D. E.; Rojc, A.; Obernier, K.; Fabius, J. M.; Soucheray, M.; Miorin, L.; Moreno, E.; Koh, C.; Tran, Q. D.; Hardy, A.; Robinot, R.; Vallet, T.; Nilsson-Payant, B. E.; Hernandez-Armenta, C.; Dunham, A.; Weigang, S.; Knerr, J.; Modak, M.; Quintero, D.; Zhou, Y.; Dugourd, A.; Valdeolivas, A.; Patil, T.; Li, Q.; Hüttenhain, R.; Cakir, M.; Muralidharan, M.; Kim, M.; Jang, G.; Tutuncuoglu, B.; Hiatt, J.; Guo, J. Z.; Xu, J.; Bouhaddou, S.; Mathy, C. J. P.; Gaulton, A.; Manners, E. J.; Félix, E.; Shi, Y.; Goff, M.; Lim, J. K.; McBride, T.; O’Neal, M. C.; Cai, Y.; Chang, J. C. J.; Broadhurst, D. J.; Klippsten, S.; De Wit, E.; Leach, A. R.; Kortemme, T.; Shoichet, B.; Ott, M.; Saez-Rodriguez, J.; tenOever, B. R.; Mullins, R. D.; Fischer, E. R.; Kochs, G.; Grosse, R.; García-Sastre, A.; Vignuzzi, M.; Johnson, J. R.; Shokat, K. M.; Swaney, D. L.; Beltrao, P.; Krogan, N. J. The Global Phosphorylation Landscape of SARS-CoV-2 Infection. Cell 2020, 182 (3), 685–712.e19. https://doi.org/10.1016/j.cell.2020.06.034.

(9) Zhu, Y.; Feng, F.; Hu, G.; Wang, Y.; Yu, Y.; Zhu, Y.; Xu, W.; Cai, X.; Sun, Z.; Han, W.; Ye, R.; Qu, D.; Ding, Q.; Huang, X.; Chen, H.; Xu, W.; Xie, Y.; Cai, Q.; Yuan, Z.; Zhang, R. A Genome-Wide CRISPR Screen Identifies Host Factors That Regulate SARS-CoV-2 Entry. Nat Commun 2021, 12 (1), 961. https://doi.org/10.1038/s41467-021-21213-4.

(10) Grodzki, M.; Bluhm, A. P.; Schaefer, M.; Tagmount, A.; Russo, M.; Sobh, A.; Rafiee, R.; Vulpe, C. D.; Karst, S. M.; Norris, M. H. Genome-Scale CRISPR Screens Identify Host Factors That Promote Human Coronavirus Infection. Genome Med 2022, 14 (1), 10. https://doi.org/10.1186/s13073-022-01013-1.

(11) Burkard, C.; Verheije, M. H.; Wicht, O.; van Kasteren, S. I.; van Kuppeveld, F. J.; Haagmans, B. L.; Pelkmans, L.; Rottier, P. J. M.; Bosch, B. J.; de Haan, C. A. M. Coronavirus Cell Entry Occurs through the Endo-/Lysosomal Pathway in a Proteolysis-Dependent Manner. PLoS Pathog 2014, 10 (11), e1004502. https://doi.org/10.1371/journal.ppat.1004502.

(12) White, J. M.; Whittaker, G. R. Fusion of Enveloped Viruses in Endosomes. Traffic 2016, 17 (6), 593–614. https://doi.org/10.1111/tra.12389.

(13) Rapiteanu, R.; Davis, L. J.; Williamson, J. C.; Timms, R. T.; Paul Luzio, J.; Lehner, P. J. A Genetic Screen Identifies a Critical Role for the WDR81-WDR91 Complex in the Trafficking and Degradation of Tetherin. Traffic 2016, 17 (8), 940–958. https://doi.org/10.1111/tra.12409.

(14) Liu, N.; Liu, K.; Yang, C. WDR91 Specifies the Endosomal Retrieval Subdomain for Retromer-Dependent Recycling. J Cell Biol 2022, 221 (12). https://doi.org/10.1083/jcb.202203013.

(15) Xing, R.; Zhou, H.; Jian, Y.; Li, L.; Wang, M.; Liu, N.; Yin, Q.; Liang, Z.; Guo, W.; Yang, C. The Rab7 Effector WDR91 Promotes Autophagy-Lysosome Degradation in Neurons by Regulating Lysosome Fusion. J Cell Biol 2021, 220 (8). https://doi.org/10.1083/jcb.202007061.

(16) Yan, B.-R.; Li, T.; Coyaud, E.; Laurent, E. M. N.; St-Germain, J.; Zhou, Y.; Kim, P. K.; Raught, B.; Brumell, J. H. C5orf51 Is a Component of the MON1-CCZ1 Complex and Controls RAB7A Localization and Stability during Mitophagy. Autophagy 2022, 18 (4), 829–840. https://doi.org/10.1080/15548627.2021.1960116.

(17) Stirnimann, C. U.; Petsalaki, E.; Russell, R. B.; Müller, C. W. WD40 Proteins Propel Cellular Networks. Trends Biochem Sci 2010, 35 (10), 565–574. https://doi.org/10.1016/j.tibs.2010.04.003.

(18) Schapira, M.; Tyers, M.; Torrent, M.; Arrowsmith, C. H. WD40 Repeat Domain Proteins: A Novel Target Class? Nat Rev Drug Discov 2017, 16 (11), 773–786. https://doi.org/10.1038/nrd.2017.179.

(19) Senisterra, G.; Wu, H.; Allali-Hassani, A.; Wasney, G. A.; Barsyte-Lovejoy, D.; Dombrovski, L.; Dong, A.; Nguyen, K. T.; Smil, D.; Bolshan, Y.; Hajian, T.; He, H.; Seitova, A.; Chau, I.; Li, F.; Poda, G.; Couture, J.-F.; Brown, P. J.; Al-Awar, R.; Schapira, M.; Arrowsmith, C. H.; Vedadi, M. Small-Molecule Inhibition of MLL Activity by Disruption of Its Interaction with WDR5. Biochem J 2013, 449 (1), 151–159. https://doi.org/10.1042/BJ20121280.

(20) Bolshan, Y.; Getlik, M.; Kuznetsova, E.; Wasney, G. A.; Hajian, T.; Poda, G.; Nguyen, K. T.; Wu, H.; Dombrovski, L.; Dong, A.; Senisterra, G.; Schapira, M.; Arrowsmith, C. H.; Brown, P. J.; Al-Awar, R.; Vedadi, M.; Smil, D. Synthesis, Optimization, and Evaluation of Novel Small Molecules as Antagonists of WDR5-MLL Interaction. ACS Med Chem Lett 2013, 4 (3), 353–357. https://doi.org/10.1021/ml300467n.

(21) Li, A. S. M.; Kimani, S.; Wilson, B.; Noureldin, M.; González-Álvarez, H.; Mamai, A.; Hoffer, L.; Guilinger, J. P.; Zhang, Y.; von Rechenberg, M.; Disch, J. S.; Mulhern, C. J.; Slakman, B. L.; Cuozzo, J. W.; Dong, A.; Poda, G.; Mohammed, M.; Saraon, P.; Mittal, M.; Modh, P.; Rathod, V.; Patel, B.; Ackloo, S.; Santhakumar, V.; Szewczyk, M. M.; Barsyte-Lovejoy, D.; Arrowsmith, C. H.; Marcellus, R.; Guié, M.-A.; Keefe, A. D.; Brown, P. J.; Halabelian, L.; Al-Awar, R.; Vedadi, M. Discovery of Nanomolar DCAF1 Small Molecule Ligands. J Med Chem 2023, 66 (7), 5041–5060. https://doi.org/10.1021/acs.jmedchem.2c02132.

(22) Kimani, S. W.; Owen, J.; Green, S. R.; Li, F.; Li, Y.; Dong, A.; Brown, P. J.; Ackloo, S.; Kuter, D.; Yang, C.; MacAskill, M.; MacKinnon, S. S.; Arrowsmith, C. H.; Schapira, M.; Shahani, V.; Halabelian, L. Discovery of a Novel DCAF1 Ligand Using a Drug-Target Interaction Prediction Model: Generalizing Machine Learning to New Drug Targets. J Chem Inf Model 2023, 63 (13), 4070–4078. https://doi.org/10.1021/acs.jcim.3c00082.

(23) McCloskey, K.; Sigel, E. A.; Kearnes, S.; Xue, L.; Tian, X.; Moccia, D.; Gikunju, D.; Bazzaz, S.; Chan, B.; Clark, M. A.; Cuozzo, J. W.; Guié, M.-A.; Guilinger, J. P.; Huguet, C.; Hupp, C. D.; Keefe, A. D.; Mulhern, C. J.; Zhang, Y.; Riley, P. Machine Learning on DNA-Encoded Libraries: A New Paradigm for Hit Finding. J Med Chem 2020, 63 (16), 8857–8866. https://doi.org/10.1021/acs.jmedchem.0c00452.

(24) Cuozzo, J. W.; Centrella, P. A.; Gikunju, D.; Habeshian, S.; Hupp, C. D.; Keefe, A. D.; Sigel, E. A.; Soutter, H. H.; Thomson, H. A.; Zhang, Y.; Clark, M. A. Discovery of a Potent BTK Inhibitor with a Novel Binding Mode by Using Parallel Selections with a DNA-Encoded Chemical Library. Chembiochem 2017, 18 (9), 864–871. https://doi.org/10.1002/cbic.201600573.

(25) Teuscher, K. B.; Chowdhury, S.; Meyers, K. M.; Tian, J.; Sai, J.; Van Meveren, M.; South, T. M.; Sensintaffar, J. L.; Rietz, T. A.; Goswami, S.; Wang, J.; Grieb, B. C.; Lorey, S. L.; Howard, G. C.; Liu, Q.; Moore, W. J.; Stott, G. M.; Tansey, W. P.; Lee, T.; Fesik, S. W. Structure-Based Discovery of Potent WD Repeat Domain 5 Inhibitors That Demonstrate Efficacy and Safety in Preclinical Animal Models. Proc Natl Acad Sci U S A 2023, 120 (1), e2211297120. https://doi.org/10.1073/pnas.2211297120.

(26) Xu, C.; Min, J. Structure and Function of WD40 Domain Proteins. Protein Cell 2011, 2 (3), 202–214. https://doi.org/10.1007/s13238-011-1018-1.

(27) Avdic, V.; Zhang, P.; Lanouette, S.; Groulx, A.; Tremblay, V.; Brunzelle, J.; Couture, J.-F. Structural and Biochemical Insights into MLL1 Core Complex Assembly. Structure 2011, 19 (1), 101–108. https://doi.org/10.1016/j.str.2010.09.022.

(28) Orlicky, S.; Tang, X.; Neduva, V.; Elowe, N.; Brown, E. D.; Sicheri, F.; Tyers, M. An Allosteric Inhibitor of Substrate Recognition by the SCF(Cdc4) Ubiquitin Ligase. Nat Biotechnol 2010, 28 (7), 733–737. https://doi.org/10.1038/nbt.1646.

(29) Han, K.; Kearnes, S.; Xu, J.; Torng, W.; Feng, J. Improving Hit-Finding: Multilabel Neural Architecture with DEL. NeurIPS 2021 AI for Science Workshop 2021.

(30) Kearnes, S.; McCloskey, K.; Berndl, M.; Pande, V.; Riley, P. Molecular Graph Convolutions: Moving beyond Fingerprints. J Comput Aided Mol Des 2016, 30 (8), 595–608. https://doi.org/10.1007/s10822-016-9938-8.

(31) Gobbi, A.; Lee, M.-L. DISE: Directed Sphere Exclusion. J Chem Inf Comput Sci 2003, 43 (1), 317–323. https://doi.org/10.1021/ci025554v.

(32) Hutchinson, A.; Seitova, A. Production of Recombinant PRMT Proteins Using the Baculovirus Expression Vector System. J Vis Exp 2021, No. 173. https://doi.org/10.3791/62510.

