## Supplemental Figures for "Discovery of a first-in-class small molecule ligand for WDR91 using DNA-encoded chemical library selection followed by machine learning"

Supplementary information

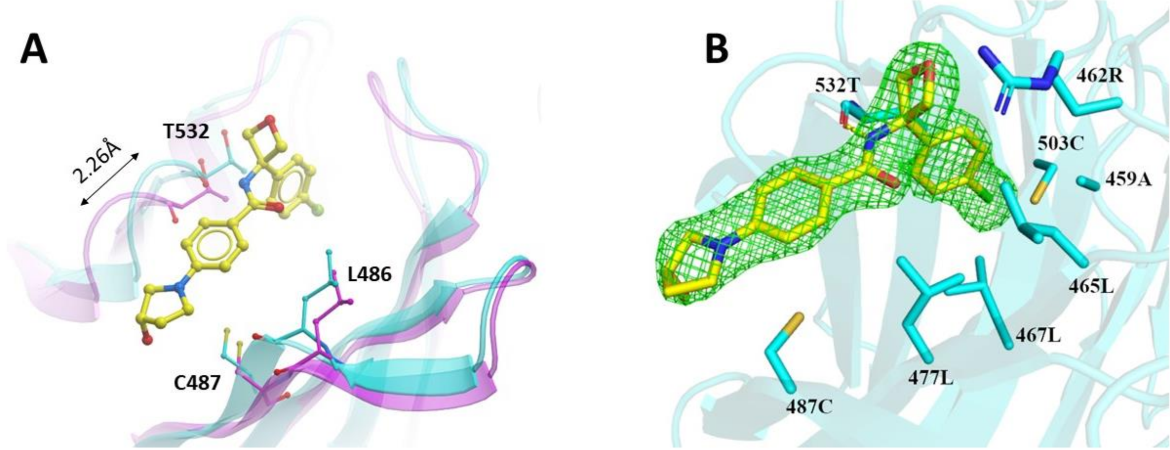


**Figure S1.** Crystal structure of WDR91-**1**. **(A)** Overlay of WDR91 apo and compound-bound structures. **(B)** Fo-Fc electron density omit map for the compound **1,** contoured at σ3 (green mesh).

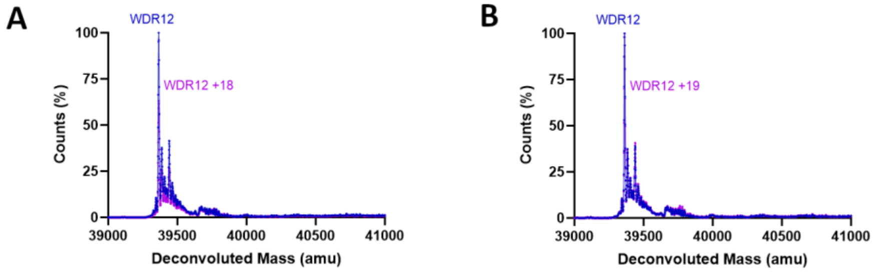


**Figure S2.** Intact mass LC/MS analysis of WDR12 (WDR) in complex with compound **18** (A) and compound **19** (B) (molar ratio of protein to ligand is 1:10). The major peak corresponds to the apo (blue) and the complex (magenta), respectively.

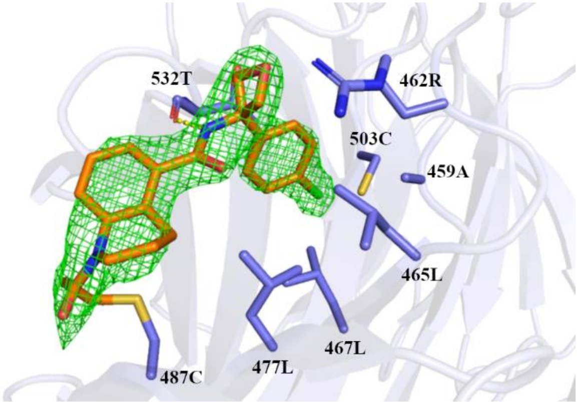

**Figure S3.** Fo-Fc electron density omit map for the compound **18** in WDR91-**18,** contoured at σ3 (green mesh).

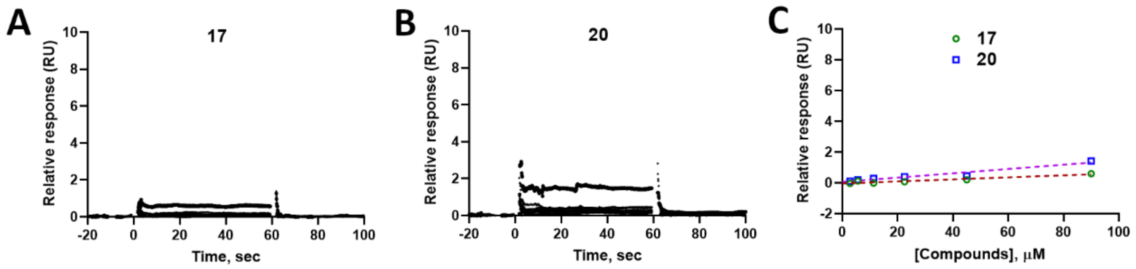


**Figure S4**. **Affinity determination by SPR**. (**A, B**) Raw sensorgrams showing no significant binding of covalent compounds **17** and **20** to ctdWDR91, and (**C**) Relative response vs concentration plots for covalent compounds **17** and **20**.

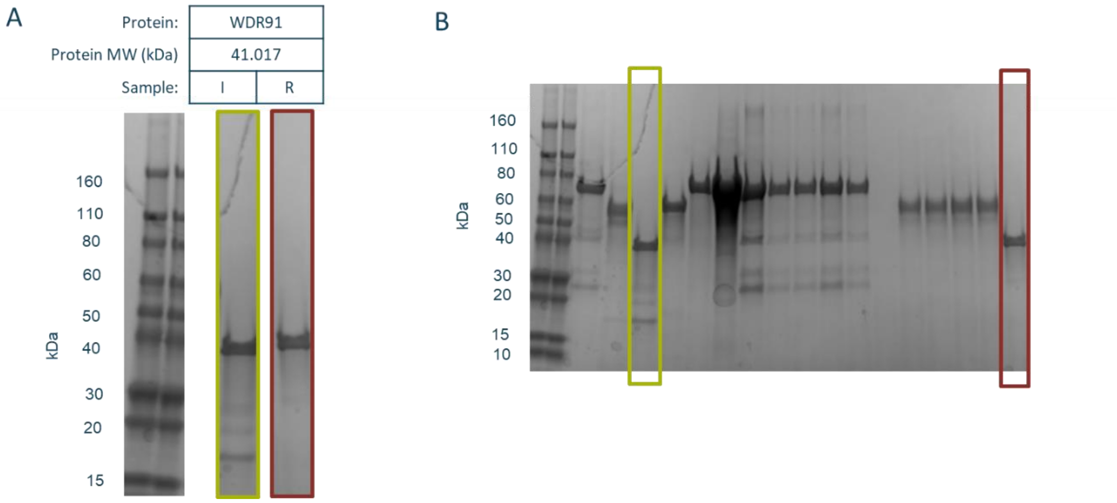


**Figure S5.** Protein Capture Assessment – A) Specific lanes of relevant samples from selections of WDR91 immobilized on IMAC resin with equivalent amounts of input (I) and resin (R) run and analyzed demonstrating >=50% of the input protein immobilized onto resin during selections for WDR91. B) Full gel containing lanes shown in panel A along with non-relevant samples.
